# Conditions for synaptic specificity during the maintenance phase of synaptic plasticity

**DOI:** 10.1101/617928

**Authors:** Marco A. Huertas, Todd Charlton Sacktor, Harel Z. Shouval

## Abstract

Activity-dependent modifications of synaptic efficacies are a cellular substrate of learning and memory. Experimental evidence shows that these modifications are synapse-specific and that the long-lasting effects are associated with the sustained increase in concentration of specific proteins like *PKMζ*. However, such proteins are likely to diffuse away from their initial synaptic location and spread out to neighboring synapses, potentially compromising synapse-specificity. In this paper we address the issue of synapse-specificity during memory maintenance. Assuming that the long-term maintenance of synaptic plasticity is accomplished by a molecular switch we perform simulations using the reaction-diffusion package in NEURON and analytical calculations to determine the limits of synaptic-specificity during maintenance. Moreover, we explore the effects of the diffusion and degradation rates of proteins and of the geometrical characteristics of dendritic spines on synapse specificty. We conclude that the necessary conditions for synaptic specificity during maintenance require that protein synthesis occurs in dendritic spines and that the activated dendritic spines exhibit small neck diameters.

## Introduction

Overwhelming experimental evidence indicates that activity-dependent modification of synaptic efficacies is the cellular substrate of learning and memory (Morris et al., 1990; Martin and Morris, 2002; Whitlock et al., 2006; Nabavi et al., 2014). Much is known about the molecular substrate of one form of synaptic plasticity, called long-term potentiation (LTP). Theoretically, such synaptic plasticity must be synapse-specific and this is supported by extensive experimental evidence (Andersen et al., 1977; Lynch et al., 1977; Harvey et al., 2008). Changes occurring at the synapse level, due to neuronal activity, include molecular alterations to the synapse machinery (Bosch et al., 2014), structural changes to dendritic spines (Matsuzaki et al., 2004; Tonnesen et al., 2014) and an increase in the synthesis of an assortment of proteins (Costa-Mattioli et al., 2009; Bosch et al., 2014). Some of these changes last for a few hours while others endure for days or a life time. A long standing problem, first raised by Crick in 1984 (Crick, 1984), is how can memories and their cellular substrate last for much longer periods of time than the life-times of the molecular substrates. A possible solution to this quandary is that local synaptic molecular switches can maintain stable synaptic efficacies even if their molecular substrates are transient (Lisman, 1985; Lisman and Zhabotinsky, 2001; Klann and Sweatt, 2008; Aslam et al., 2009; Jalil et al., 2015).

Over the last couple of decades an accumulation of evidence has shown that a sustained increase in the concentration of specific isoforms of *PKC* is associated with the long-lasting form of LTP (L-LTP) (Tsokas et al., 2016; Wang et al., 2016). Chiefly among these *PKMζ*, an atypical isoform of *PKC*, has been shown to be necessary and sufficient for some forms of long-term plasticity and memory (Sacktor et al., 1993; Drier et al., 2002; Ling et al., 2002; Wang et al., 2016; Tsokas et al., 2016). However, in mutants lacking the gene for *PKMζ* another isoform of atypical *PKC, PKCι/λ*, is upregulated and becomes necessary for maintenance. Experimental evidence indicates that polyribosomes, responsible for protein synthesis, exist both in dendrites and spines, and that the induction of L-LTP increases the number of them in synaptic spines (Sutton and Schuman, 2006; Bourne et al., 2007). Therefore, the likely substrate for the molecular switch that maintains memory and, at the same time, is synapse specific is implemented at the level of the synthesis of new proteins (Aslam et al., 2009), and most likely the synthesis of PKMζ (Jalil et al., 2015).

Thus far, studies of bi-stable switches as the basis for the maintenance phase of synaptic plasticity have concentrated on a single compartment, and have shown that switches based on positive molecular feedback can be bi-stable or multi stable. However, if these switches are located within dendritic shafts or dendritic spines, their products can diffuse and affect other switches in their vicinity. Such interaction could potentially eliminate synapse-specificity during maintenance. Moreover, considering that *PKMζ* might be degraded slowly (van de Nes, 2012; Wang et al., 2016) and is constitutively active due to the lack of the N-terminal regulatory domain (reviewed in van de Nes (2012)), this can result in a long protein length-constant, as explained below. Such a long length-constant means that switches at distant locations can potentially interact and place a very severe constraint on synapse-specificity.

In this paper we address the issue of synapse-specificity during maintenance, employing a computational approach based on reaction-diffusion equations in dendritic shafts and spines. We study this equation using both an analytical approach and the reaction-diffusion package of the NEURON simulation platform (McDougal et al., 2013; Carnevale and Hines, 2006). This analysis allowed us to estimate the limits of synapse-specificity when the molecular switches are located in dendritic shafts or in dendritic spines and how these limits depend of the system parameters. These results allow us to make strong predictions that can be tested experientially.

## Materials & Methods

We developed a spatial model consisting of a dendritic branch with a variable number of dendritic spines, and numerically calculated the steady-state spatial distribution of proteins being synthesized at various locations in the model’s volume. Proteins are synthesized in specified locations in the dendrite or in the head of dendritic spines. Furthermore, proteins can be degraded anywhere and are allowed to diffuse throughout the defined volume. Here we describe the various elements and setup of the model.

### Protein synthesis model

Protein synthesis by polyribosomes is modeled using a bi-stable switch that includes a concentration-dependent synthesis and degradation components. The synthesis rate is depended on the protein concentration at the location of the polyribosome (Kelly et al., 2007; Costa-Mattioli et al., 2009), while the degradation rate depends on the local protein concentration throughout the volume. In general, this non-linear positive feedback can generate bi- or multi-stability solutions (Lisman, 1985; Lisman and Zhabotinsky, 2001; Aslam et al., 2009; Jalil et al., 2015). In the absence of diffusion, the change in protein concentration (c), at one point, is described by the equation

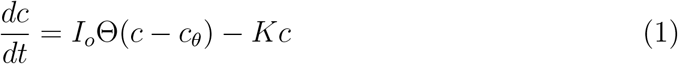

where *I_o_* is the maximum protein synthesis rate, *c_θ_* is an activation threshold and *K* is the protein degradation rate. The activation function Θ is assumed to have a general sigmoid shape. In the analytical calculations presented here we will assume that Θ has a step-function form.

The value of *I_o_* is chosen to guarantee a bi-stable steady-state solution. Figure 1A shows the fixed points of Eq. 1 (i.e., when *dc/dt* = 0) for different values of *I_o_*, while keeping the degradation rate *K* constant. As illustrated here, for values of *I_o_* below 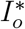 there is only one fixed point at *c* = 0. This solution will be referred to as down- or inactive-state. For values above 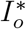 there is another stable solution at higher concentrations. This corresponds to the up- or active-state.

**Figure 1:**
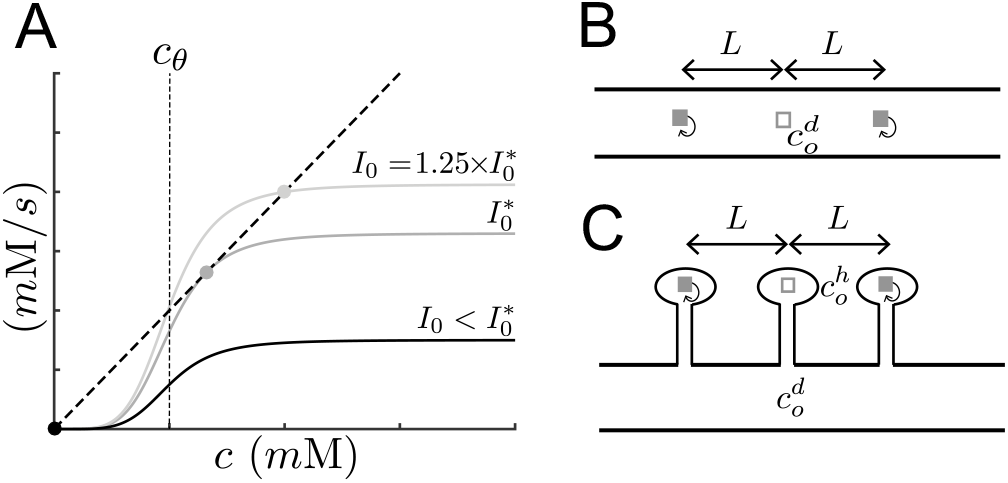
Bi-stable switch and model setup. A) Fixed points of Eq. 1 for various values of *I_o_*. Below 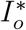 there is one fixed point at *c* = 0 (down-state), whereas for values above 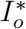 there is a second fixed point at larger concentrations (up-state). The value of 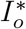 is the smallest value of *I_o_* required to produce an up-state. The activation curve Θ illustrated here has a sigmoid dependency in protein concentration. B) Schematic of the case of polyribosomes in the dendrite. Two active polyribosomes (gray boxes with feedback arrows) are located a distance *L* on each side of a third inactive polyribosome (empty white box). The protein concentration at the location of the inactive polyribosome is labeled 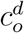. C) Similar as in B for the case of polyribosomes in dendritic spine heads. The protein concentration at the location of the inactive polyribosome is labeled 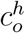, whereas the concentration on the dendrite at the location of the spine is 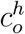.

We consider that for active synapses the switch is operating in the up-state of the bistable regime. In the case when Θ is represented by a step-function, the up-state lies in the saturation region, otherwise if it has a sigmoidal shape the up-state will be just below saturation. Therefore, we use 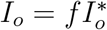, with *f* > 1. In simulations we chose *f* = 1.25. Clearly the value of 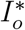 will be different for the various conditions investigated here.

### Model setup: multiple polyribosomes

To determine the limits of the ability of synapses to maintain their specificity during the maintenance phase, we consider the following setup. We place 2*N* + 1 polyribosomes (i.e., bi-stable switches) equidistant from each other along the length of a dendritic branch. One polyribosome is located at *x* = 0 and is considered to be in the down-state while the remaining ones, separated a distance *L*, are in the upstate presumably as the result of synapses being potentiated. Figure 1B shows this situation for the case *N* = 1.

Proteins synthesized by the active polyribosomes diffuse along the dendrite and can cause an increase in the concentration at the location of the inactive polyribosome. The concentration at this location will depend on the distance *L* between them and the number of active polyribosomes. We determine the smallest distance (*L_crit_*) at which the inactive polyribosome remains in the down-state when *N* → ∞. The *N* → ∞ is the most stringent limit, and additionally the results presented are then simpler as they are independent of this limit. However, the results do not strongly depend on this limit, and using a moderate value of *N*, (¿5) will cause only a small diffrence in the results. The value of *L_crit_*, which we refer as the critical distance, provides a measure of the distance between activated synapses that will keep an inactive synapse isolated.

A similar setup is made for the case of polyribosomes located in the heads of dendritic spines, as illustrated in Fig. 1C. Here the distance *L* refers to the inter-spine distance and thus *L_crit_* measures how close dendritic spines (and their corresponding synapses) could be from each other and still remain isolated.

### Analytical expressions for *L_crit_*

The value of *L_crit_* can be calculated analytically. In the case of polyribosomes along a dendrite we assume that all active polyribosomes are operating in the saturation regime with a maximum synthesis rate *I_o_*. In this case the concentration at the location of the inactive polyribosome takes the form

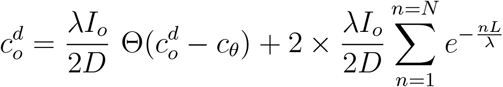

where *λI_o_*/2*D* is the maximum concentration a the location of an isolated polyribosome (see Eq. 6). The first term in this expression represents the contribution from the inactive polyribosome, whereas the second term is the contributions of the 2*N* active polyribosomes. The exponential function arises from the solution to the one-dimensional diffusion equation with a concentration-dependent degradation rate (cf. Eq. 6). The geometric series has the closed form, resulting in the equation:

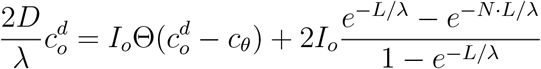

In the limit of *N* → ∞ this expression becomes:

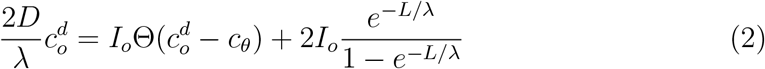

Note, that the difference between the two equations above is the term *e^−N·L/λ^*. When polyribosomes reside in dendrites, this term is very small even for moderates values of *N*. For example a solution for *N* = 5, differs from the infinite limit by a few percent. We solve this equation for 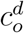 —the concentration at the location of the inactive polyribosome. The solutions of Eq. 2 for various values of *L* and a step function Θ are graphically illustrated in Fig. 2. As L becomes smaller the solutions undergo a transition from being bi-stable to mono-stable. The value at which this transition occurs defines *L_crit_* because the only solution possible at smaller distances will render all polyribosomes in the up-state.

**Figure 2:**
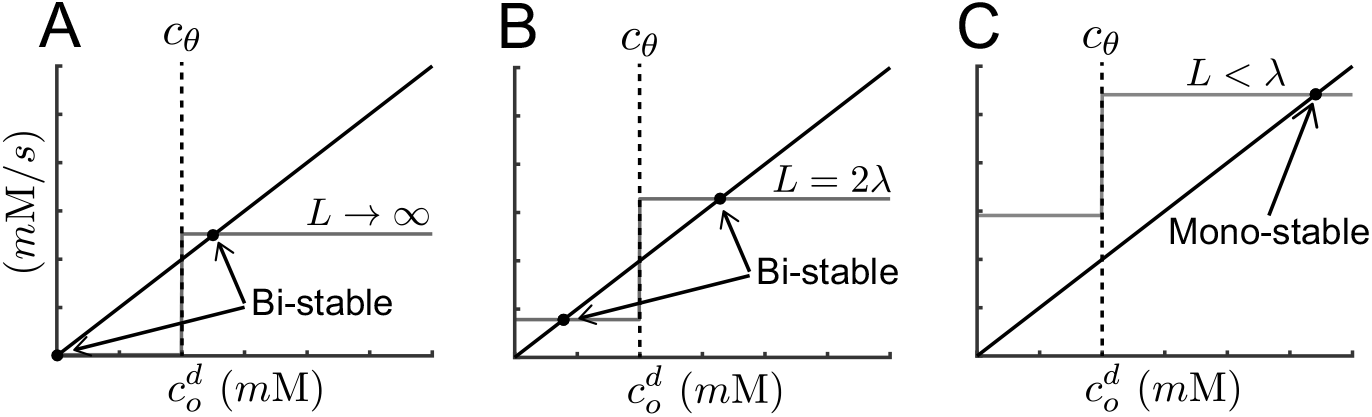
Transition from bi-to mono-stability as a function of inter-polyribosome distance for the case of an infinite number of sources in the dendrite. Panels A, B and C illustrate the effect of the distance between polyribosomes on the number of stable fixed points in Eq. 2. Fixed points are indicated by black circles and correspond to the intersection between the activation curve (gray step-function) and decay rate curve (solid black line). The three examples correspond to different distances between polyribosomes: A) *L* → ∞, B) *L* = 2λ and C) *L* < λ. The system undergoes a transition from bi-stability (*L* > λ) to mono-stability (*L* ≤ λ).

A similar analysis can be done when polyribosomes are located in the head of dendritic spines. However in this case we are interested in the concentration in the spine head 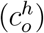. Due to diffusion through the dendritic spines the concentration *c^h^* depends on both *I_o_* and 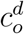, the concentration in the dendrite at the location of the spine (see Fig. 1C)

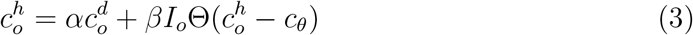

where *α* and *β* are parameters determined by the spine geometry, the diffusion constant D and the characteristic length constant λ (see section Parameters and auxiliary functions). The label *o* serves to identify these quantities as pertaining to the spine at *x* = 0. The value of 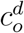 is determined by superposing the contributions from the inactive spine and the contributions from the remaining 2*N* spines. The final expression becomes, in the limit of *N* → ∞

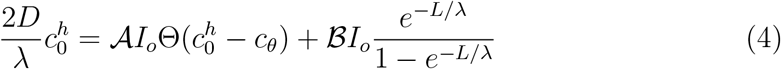

where 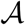 and 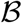 are constant functions of the geometry of the spines, the diffusion constant *D* and the length constant λ (see section Parameters and auxiliary functions). This equation resembles Eq. 2 above and thus we can determine the critical value *L_crit_* similarly, as depicted in Fig. 2.

Equations 2 and 4 are the main equations used for this study.

### Solutions for isolated polyribosomes

Expressions Eq. 2 and 4 depend on the maximum synthesis rate *I_o_*. This value is determined from the condition that the isolated polyribosome remains in the upstate against the effects of diffusion and degradation (cf. Fig. 1A). In the case of a polyribosome in the dendrite, the relevant expression is derived from the steady-state solution to the one-dimensional reaction-diffusion equation

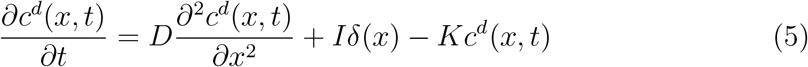

where the label ^*d*^ indicates the concentration is in the dendrite, 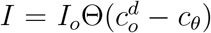 is the synthesis rate with a maximum value *I_o_* and 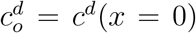. The solution is given by

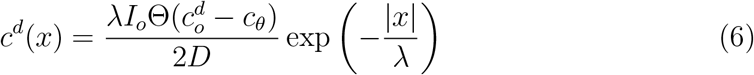

where 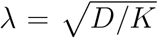 is the characteristic length constant of the protein. Evaluating this solution at x = 0, provides an expression from which to determine the value of 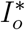:

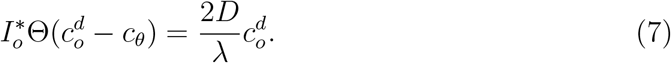

This equation is similar to the fixed point of Eq. 1 with *K* replaced by 2*D*/λ and thus, as illustrated in Fig. 1A, the value of 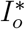 is determined so that 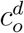 is in the up-state.

In the case of polyribosomes in dendritic spines we first solved Eq. 5 in each compartment representing the dendritic spine, see Fig. 3. Solutions for each compartment are matched at the boundaries. Moreover the fluxes between compartments are related through the expression 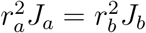, where 2*r* is the diameter and *J* is the flux in or out of the compartment. Here we have used a simplified expression that agrees with the simulations. A more elaborated formulation can be found in Berezhkovskii et al. (2009) however, for our conditions the expressions are identical.

**Figure 3:**
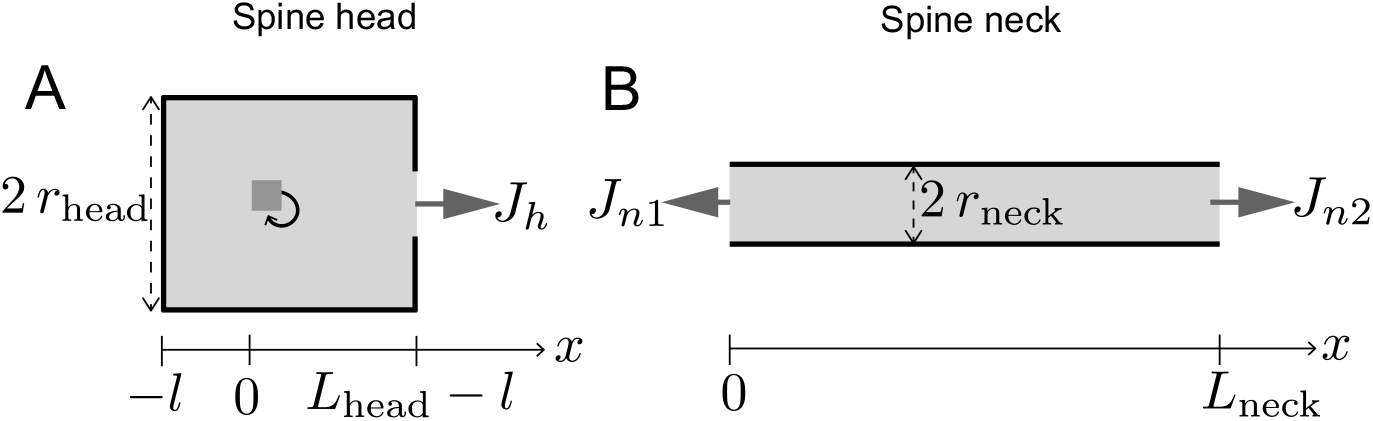
Geometry of spine. Panels A and B show schematics of the compartments representing the spine head and neck, respectively. Compartments are assumed to be cylindrical in shape with diameters 2*r*_head_ and 2*r*_neck_, and lengths *L_head_* and *L_neck_*, respectively. The polyribosome (gray box with a feedback arrow) is located a distance *l* from the sealed end of the spine head compartment. The quantities *J_h_*, *J*_*n*1_ and *J*_*n*2_ represent protein fluxes per unit area out and in of the respective compartments.

The resulting equations are combined to derive the expression

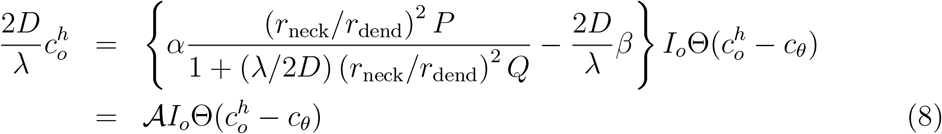

where 2*r*_neck/dend_ is the diameter of the neck/dendritic compartment and *α*, *β*, *P* and *Q* are terms containing geometrical, diffusion and degradation parameters (see section Parameters and auxiliary functions). This equation is used to find 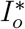 under the same conditions as those for Eq. 7, i.e., to guarantee that the solution corresponds to the up-state.

### Computational model and simulations

Simulations were performed using the reaction-diffusion package (McDougal et al., 2013) in NEURON (Carnevale and Hines, 2006) to solve for the steady-state concentration distribution of protein. Proteins were allowed to diffuse and degrade in a volume representing a dendritic branch with a variable number of dendritc spine compartments. The dendritic branch was considered as a cylinder of diameter ddend and length *L*_dend_, and the spine geometry is illustrated in Fig. 3. All the code is written in Python. The parameters used in the simulations performed are given in Table 1. The characteristic length constant, λ (see Eq. 7), which depends both on the diffusion and the degradation rate, is explored as a variable parameter in this model. The length of the dendritic branch was adjusted depending on the value of λ to guarantee that the concentration was exactly zero at the edges of the compartment. The number of segments were chosen to give a spatial grid of 1*μ*m. In the simulations the activation curve Θ was modeled as a steep Hill function,

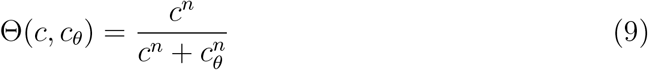

**Table 1:** Model parameters used in simulations. ^†^The expression for λ is give below Eq. 6 in terms of the degradation constant *K*.

| Model parameters |  |  |  |
| --- | --- | --- | --- |
| Diffusion coefficient | $D$ | $1 \times 10^{-3}$ | $\mu\text{m}^2/\text{ms}$ |
| Characteristic length constant | $\lambda^\dagger$ | variable | $\mu\text{m}$ |
| Hill function exponent | $n$ | 40 | — |
| Activation threshold | $c_\theta$ | 2 | mM |
| Diameter dendritic branch | $2r_{\text{dend}}$ | 5 | $\mu\text{m}$ |
| Length dendritic branch | $L_{\text{dend}}$ | variable | $\mu\text{m}$ |
| No. segments dendritic branch | $N_{\text{dend}}$ | variable | — |
| Diameter spine head | $2r_{\text{head}}$ | 1 | $\mu\text{m}$ |
| Length spine head | $L_{\text{head}}$ | 1 | $\mu\text{m}$ |
| No. segments spine head | $N_{\text{head}}$ | 5 | — |
| Diameter spine neck | $2r_{\text{neck}}$ | 0.2 | $\mu\text{m}$ |
| Length spine neck | $L_{\text{neck}}$ | 5 | $\mu\text{m}$ |
| No. segments spine neck | $N_{\text{neck}}$ | 25 | — |

Simulations that approximate the limiting case of an infinite number of polyribosomesin dendrites or spines (see Figs. 5A, 7A and 9) were performed using a finite number of sources. The number was chosen so that no significant change was observed in the concentration at the location of the inactive synapse when an additional pair of sources was added. This number was different for different values of λ. Typical values were between 10 to 30.

### Parameters and auxiliary functions

Model parameters are presented in Table 1 and the terms involving geometrical, difussional and degradation parameters in Eqs. 3, 4 and 8 are given here.

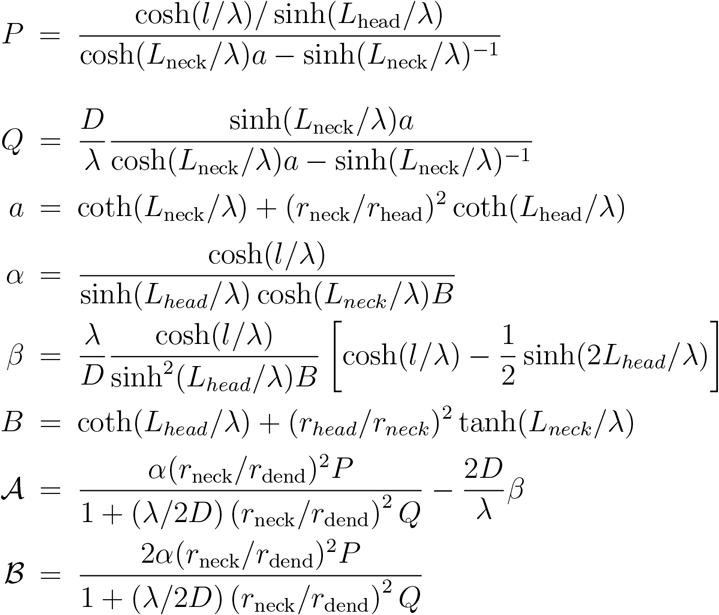

## Results

In order to maintain memory and the ability to persistently perform learned tasks over long periods of time, synaptic plasticity must generate stable changes in synaptic efficacies. How this is done is not obvious since the changes in protein number and function are typically transient due to protein turnover and diffusion. One way to obtain such long time scales is through bi- or multi-stable molecular switches (Lisman, 1985; Lisman and Zhabotinsky, 2001; Aslam et al., 2009; Jalil et al., 2015). We have argued that such a switch is most likely implemented at the level of translation, and that the synthesized protein that potentiates synaptic transmission is PKMζ. The approach we take is based on the observation that its synthesis is local in dendrites and perhaps spines (Muslimov et al., 2004; Westmark et al., 2010; Palida et al., 2015). The synthesis of a specific protein (for example PKMζ) could affect its own level of translation, and this positive feedback loop can generate bi or multi-stable switches (Westmark et al., 2010). Much of our results here though are general and could apply to other forms of a molecular switch.

A molecular switch for maintaining synaptic efficacies, must operate in a synapse-specific manner in order to maintain the computational power of the neural circuits. Until now most models of such a molecular switch were single compartment models that did not analyze the effect of diffusion on synapse-specificity. Diffusion, however, could impair synapse-specificity because proteins synthesized by one polyribosome could diffuse to a neighboring one and trigger protein synthesis at that location too. It is also thought that maintenance-related proteins have long lifetimes. Such long lifetimes, as shown below, are actually potentially detrimental to synapse-specificity.

Using the model outlined above of bistable switches in dendritic shafts or spines we have calculated the conditions necessary for synapse-specificity during the maintenance phase of synaptic plasticity.

### Polyribosomes in dendrites

To determine when a synaptic site remains isolated from neighboring potentiated synapses we consider the situation of an inactive polyribosome flanked by 2*N* active polyribosomes separated from each other by a distance *L* (see Materials & Methods). This situation is illustrated in Fig. 4A for the case of *N* = 2.

**Figure 4:**
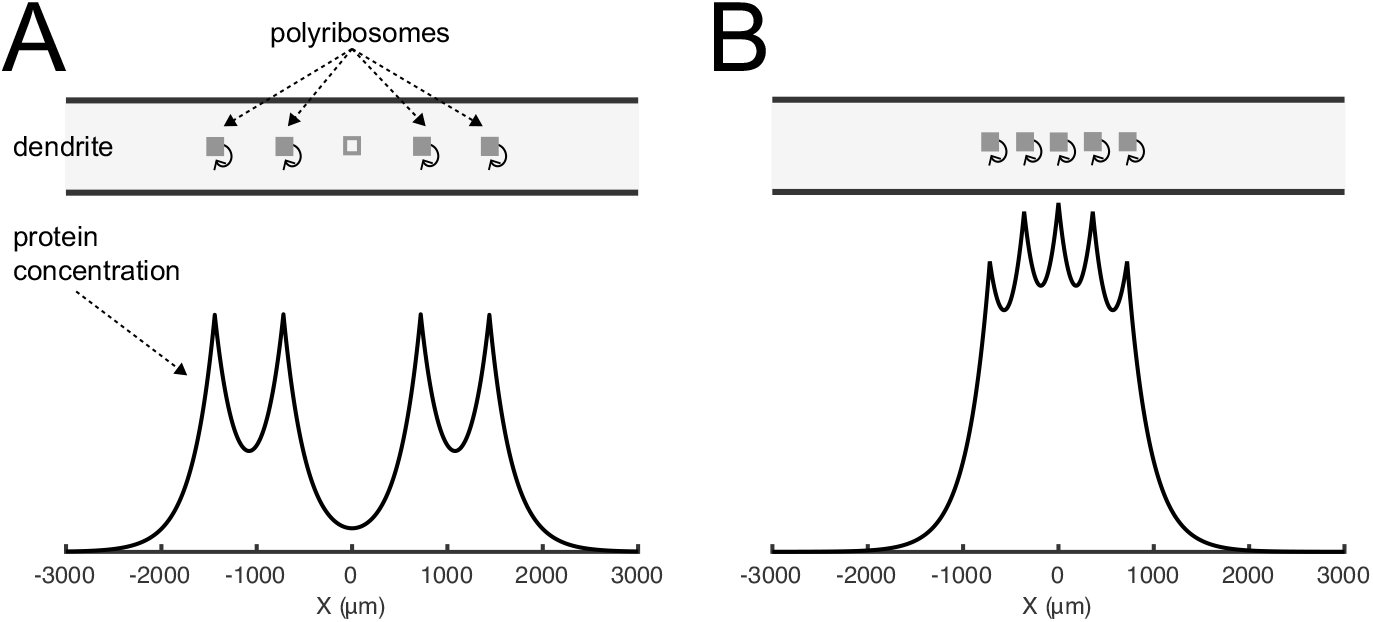
Transition between up and down states for polyribosomes located in dendrites. A) Upper panel illustrates an arrangement of active polyribosomes (gray filled boxes) on both sides of an inactive one (open box) along a dendrite. Lower panel shows a steady-state protein concentration profile. The concentration peaks at the location of the active polyribosomes. B) Similar as in A but with polyribosomes located closer to each other. Here the central polyribosome has become active.

As described above synthesized proteins can diffuse to neighboring sites leading to an increase in protein concentration at the location of the inactive polyribosome, which could result in its activation. The lower panel of Fig. 4A shows the calculated steady-state spatial profile of protein concentration along the length of the dendritic shaft—computed using NEURON. As expected the protein concentration peaks at the location of the active polyribosomes. In the case illustrated here the distance separating the polyribosomes is such that the protein concentration at the site of the inactive polyribosome (x = 0) is low enough that it does not lead to its activation.

However the situation changes when the distance between the polyribosomes decreases. As illustrated in Fig. 4B the distance between polyribosomes is short enough that the inactive polyribosome becomes active. In this example an activation pattern (e.g., memory) that involved only the four neighboring synapses has now been modified by the effects of diffusion, loosing its synaptic specificity.

As described in Materials & Methods as the distance L decreases, the number of fixed points of Eq. 2 change from two to one. Figure 5A shows a plot of the value of the stable (black solid lines) and unstable (black dashed lines) fixed points as a function of the distance L. The vertical red dashed line indicates the distance at which a transition occurs from a bistable to a monostable region. This line defines the quantity *L_crit_*.

**Figure 5:**
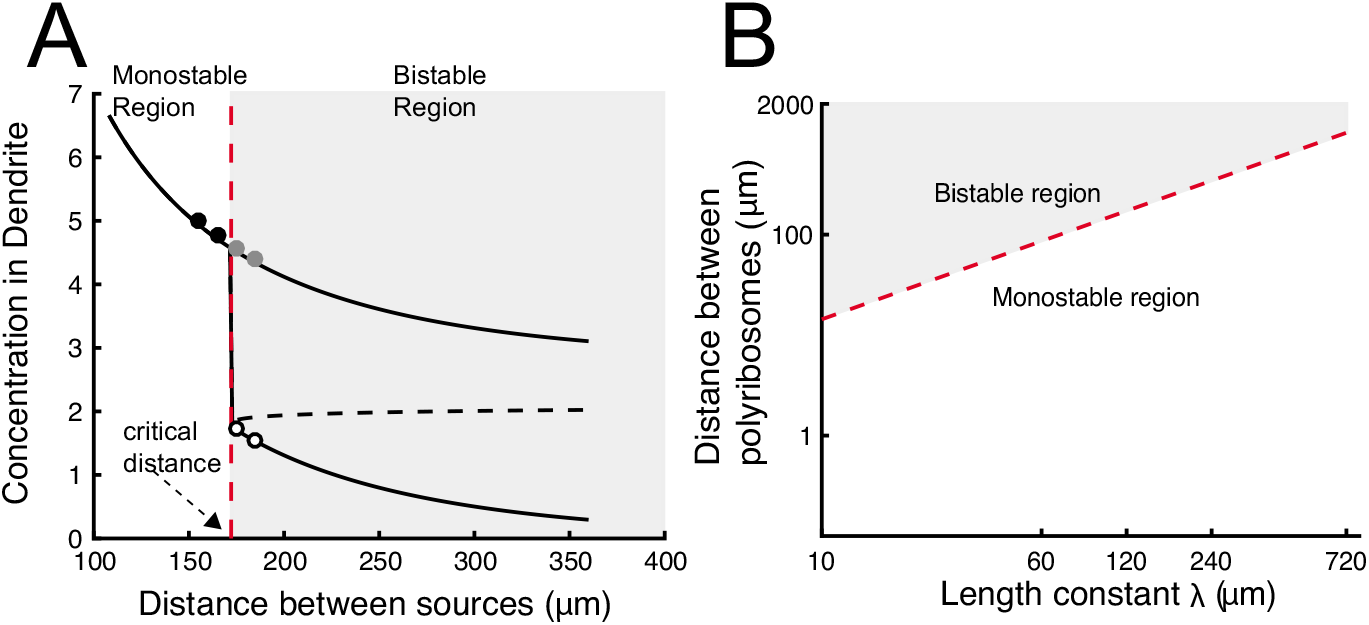
Transition of polyribosome state when polyribosomes are distributed in the dendrite. A) Example showing the transition from a bi-stable to a mono-stable mode for the central polyribosome illustrated in Fig. 4A as the distance between polyribosomes decreases. Black solid and dashed lines represent stable and unstable fix points of Eq. 2, respectively as a function of polyribosome separation. Circles correspond to simulations performed near the transition (vertical red dashed line). Filled symbols correspond to up-state and open circles to down-state solutions. B) Diagram illustrating the dependency of critical distance as a function of the characteristic length constant λ.

Figure 5 shows that for distances larger than *L_crit_* there are two stable fixed points. The upper branch of the bistable region corresponds to the situation when all polyribosomes are active. In contrast the lower branch represents the situation described in which a synapse can stay weak even though all its neighbours are potentiated. When the distance is smaller than *L_crit_* the only stable solution is for all polyribosomes to be active, thus there is a loss of synaptic specificity.

The value of *L_crit_* depends on several parameters, as can be inferred from Eq. 2. However we will focus here on the effect of the characteristic length constant, λ, since this incorporates the effects of both diffusion and degradation (defined in Materials & Methods as 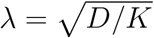). Figure 5B shows this dependence. The red dashed-line separates the mono- and bi-stable regions and corresponds to the value of *L_crit_* for different values of λ.

In general we see that with polyribosomes in the dendrite the characteristic value of *L_crit_* (dashed red line) is on the order of 10^2^*μ*m for proteins that degrade slowly. For example using a diffusion coefficient *D* = 10^-3^*μ*m^2^/ms, typical for proteins the size of PKMζ, and proteins degrading with a time constant of ~ 5 h, the corresponding value of λ is 120 μm. In this case the critical distance is on the order of 160 *μ*m which is hundreds of times larger than estimated inter-spine distances.

An analytical expression for this dependency can be given in the case of Θ being a step-function. By using Eq. 2 in the case when *c_o_* < *c_θ_*, and using the expression for the maximum synthesis rate 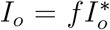 (see Materials & Methods). The value of 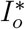 is calculated by Eq. 7 to obtain the relationship 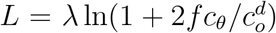. Because *L_crit_* occurs when 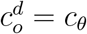 and using the value *f* = 1.25, we obtain

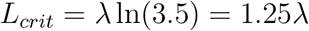

### Polyribosomes in spine heads

A similar analysis can be made when polyribosomes are located in the head of dendritic spines. In this case the arrangement of the spines is illustrated in Fig. 6A. Here an inactive polyribosome is placed in the head of a spine situated at the origin. This spine is flanked by 2*N* other spines with active polyribosomes in their corresponding heads.

**Figure 6:**
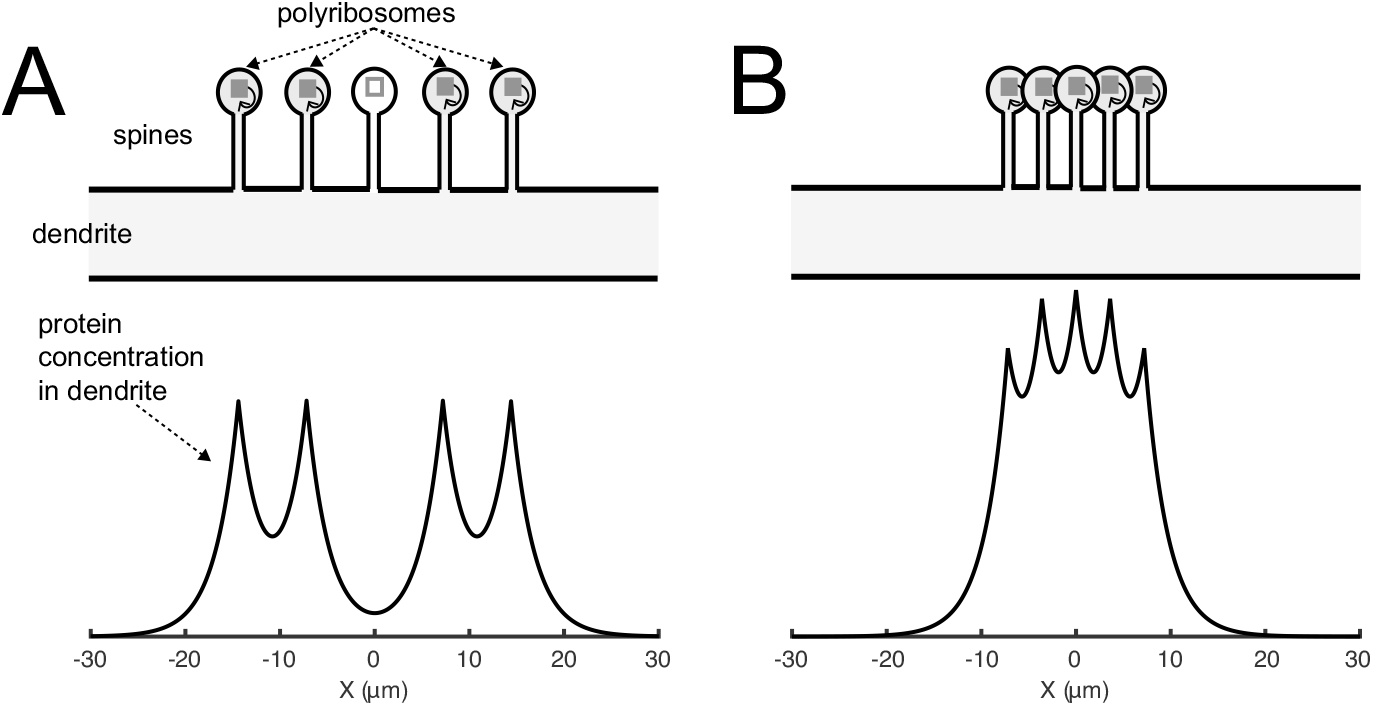
Transition between up and down states for polyribosomes located in the head of dendritic spines. A) Upper panel illustrates an arrangement of active polyribosomes(gray filled boxes) on both sides of an inactive one (open box), all located in the heads of dendritic spines distributed along a dendrite. Lower panel shows the steady-state protein concentration profile as measured in the dendrite. The concentration peaks at the location of the spines with active polyribosomes. B) Similar as in A but with spines placed closer to each other. In this situation the polyribosome in the central spine has become active. Compare with Fig. 4.

Proteins synthesized in the spine head diffuse through the spine necks into the dendrite where they can reach the inactive polyribosome. The lower panel of Fig. 6A shows the calculated steady-state protein concentration profile resulting in the dendrite. Note that the scale in the axis has decreased by a factor of 100 compared with Fig. 5A.

In a similar way, as described in the previous section, as the spines get closer to each other the concentration in the head of the inactive polyribosome increases and at the critical distance it will cause the polyribosome to become active. This is illustrated in Fig. 7B. As described in Materials & Methods the number of stable fixed-points of Eq. 4 changes as the distance L between spines decreases. The distance at which this transition occurs corresponds to *L_crit_*. An example is shown in Fig. 7A.

**Figure 7:**
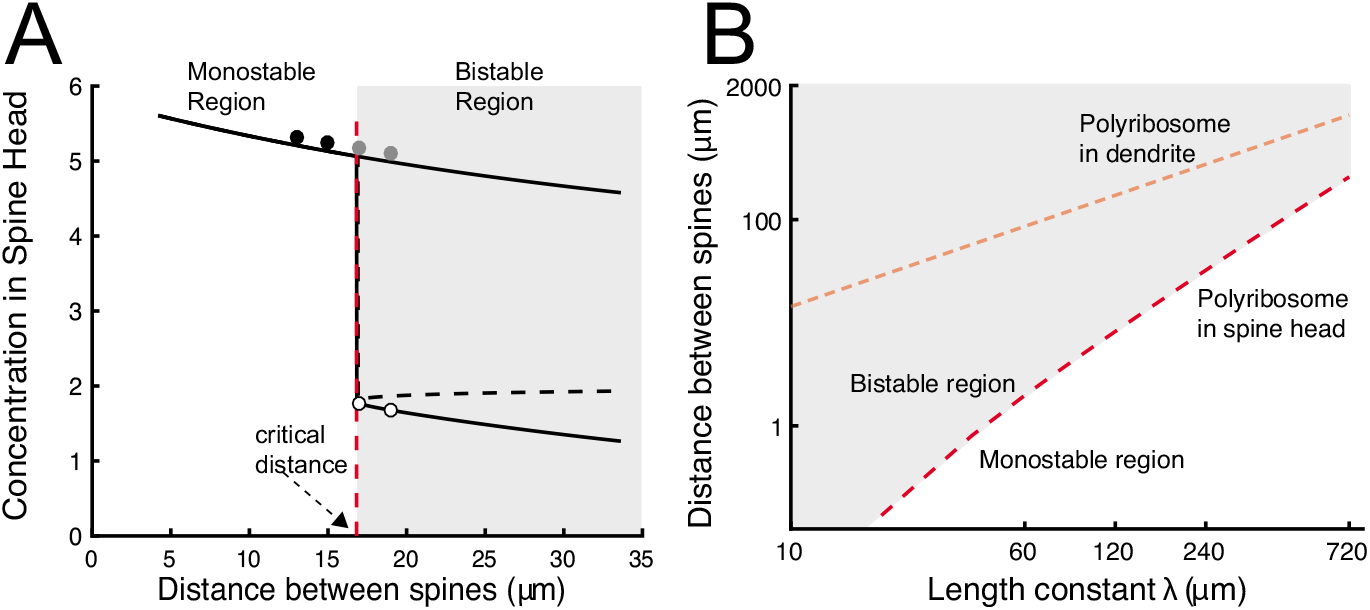
Transition of polyribosome state when polyribosomes are in located in the heads of dendritic spines. A) Example showing the transition from a bi-stable to a mono-stable mode for the polyribosome in the central spine (cf. Fig. 6A) as the distance between spines decreases. Black solid and dashed lines represent stable and unstable fix points of Eq. 4 as a function of spine separation. Circles correspond to simulations performed near the transition (vertical red dashed line). Filled symbols correspond to up-state and open circles to down-state solutions. B) Diagram illustrating the dependency of critical distance as a function of the characteristic length constant λ. Red dashed curve represents the critical distance for polyribosomes in spine heads. Light red dashed curve is the result when polyribosomes are in the dendrite (cf. Fig. 5B).

As before, as the distance between spines decreases the protein concentration in the spine head of the inactive spine also increases; however, at a distance *L_crit_* there is an abrupt change in the protein concentration. Below this distance the system can operate in a bistable region, therefore it is possible to have all spines active except for the one at x = 0. However when the distance is smaller than Lcrit there is only one possible stable fixed point, i.e., all spines are active. In this plot we see that *L_crit_* is slightly larger than 15 *μm*. This result is an order of magnitude smaller than what is obtained when the switches are located in the dendritic shaft. These analytical results were confirmed by numerical simulations (circles).

The value of *L_crit_* depends on the model parameters, for example on λ as shown in Fig 7B. This plot also shows a comparison of the critical distance between spines for various values of λ and the corresponding distance for polyribosomes located in the dendrite. Note that the axes are logarithmically scaled.

Clearly, for the same characteristic length constant, when polyribosomes are located in the head of dendritic spines the distance between these spines —that provide synaptic specificity— is significantly smaller than that obtained when polyribosomes are located in dendritic shafts. As λ increases so does the critical distance and for values of λ ~ 720*μ*m the distance is much larger than charectaristic inter-spine distances. Typical spine density in neurons has been reported to be approximately ~1 spines/*μ*m (Kirov et al., 1999; Sala and Segal, 2014). One must note that there are different spine types, and not all of them form functional synapses; this estimate groups different synaptic types together.

An analytical expression for *L_crit_* can be derived under the assumption of a step-function activation:

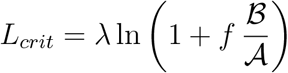

The finctions 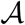 and 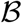, given in section Materials & Methods, both depend on λ. Therefore, this relationship is non-linear in λ.

### Dependence of critical inter-spine distance on diffusion rate and spine morphology for polyribosomes in dendritic spine heads

The critical distance between spines, *L_crit_*, can be affected by various properties of the dendritic spines as can be inferred from Eq. 4 in Materials & Methods. Here we explored the effects of having a slower diffusion coefficient in the spine than the dendrite and the influence of various morphological parameters. Note that changes in these parameters will affect the ability of an isolated active spine to remain in the up-state; therefore we recalculated the magnitude of the maximum synthesis rate 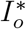 for each data point (see Eq. 8).

The presence of actin in the cytoskeleton (Matus, 2000; Shirao and Gonzalez-Billault, 2013) of the spine can slow the diffusion of proteins compared to conditions in the dendrite, decreasing effectively the diffusion coefficient inside the spine. We reasoned that such a decrease should in principle affect both the flux out and into the spine neck and could cause the inactive polyribosome to remain so unless the spines get much closer, leading to a smaller value of *L_crit_*. However, as illustrated in Fig. 8A the value of *L_crit_* does not change significantly as the ratio *D*_spine_/*D*_dendrite_ decreases, for constant *D*_dendrite_. Even when *D*_spine_ is only 1% of *D*_dendrite_ the reduction of *L*_crit_ is minimal for the values of λ considered here (λ is calculated using *D*_dendrite_).

**Figure 8:**
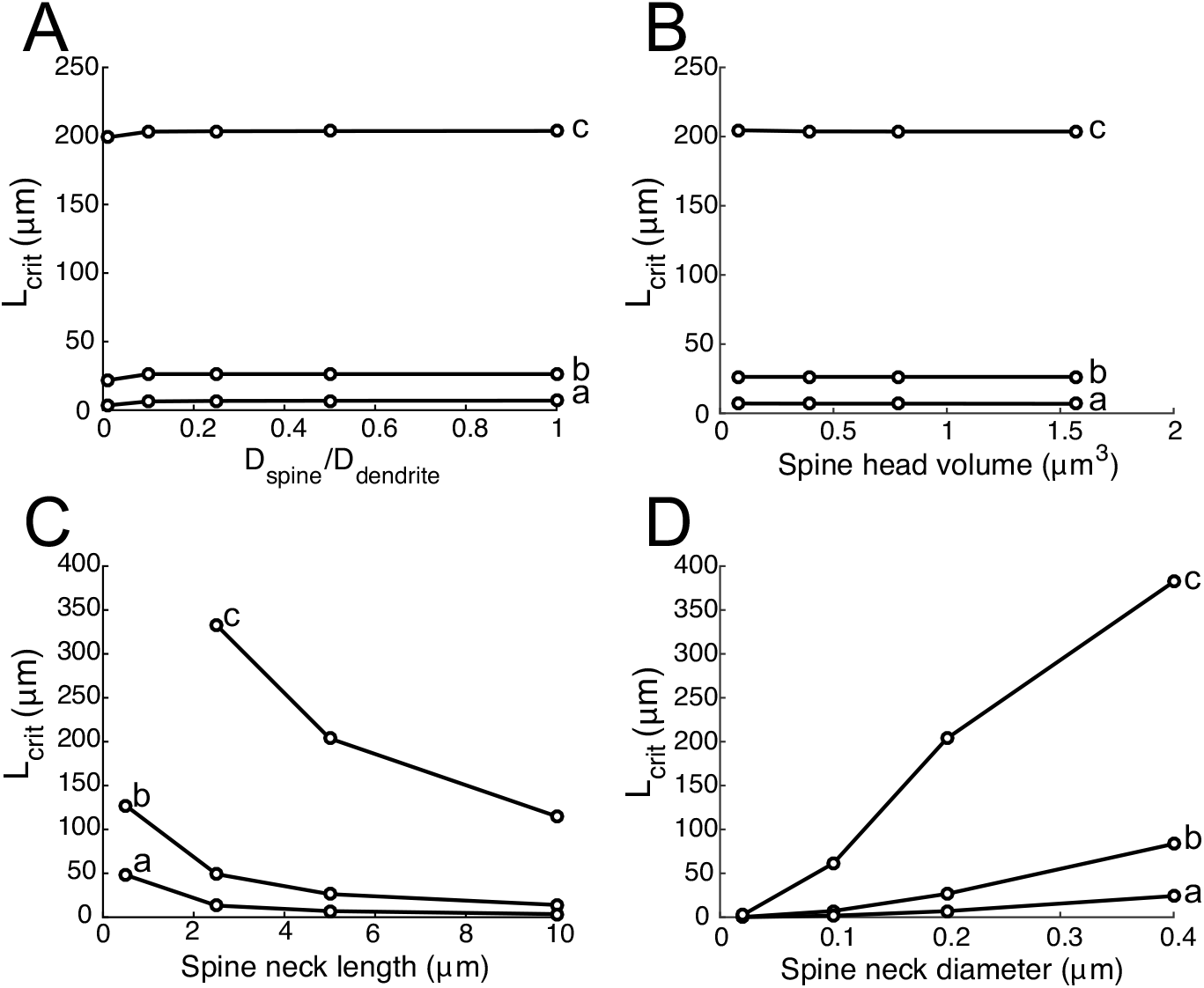
Dependency of *L_crit_* on spine morphology. Panels A-D illustrate how the minimum distance between spines depends on various parameters describing the dendritic spine. The three traces labeld a, b and c correspond to different values of the characteristic length constant λ, respectively 120, 240 and 720 *μ*m. The parameters are: A) diffusion coefficient in spine, B) spine head volume, C) spine neck length and D) spine neck diameter.

The morphology of the spine head and neck can also play a role in *L_crit_*. There are numerous reports showing a persistent change in the volume of spine heads in potentiated spines (Lee et al., 2009; Govindarajan et al., 2011; Tonnesen et al., 2014). We explored here the effects on Lcrit for various head sizes. Our results show that changes in the spine head’s volume have no effect in the critical distance as shown in Fig. 8B.

In addition to the above morphological changes, there is evidence of changes in spine’s neck length (Araya et al., 2014). Although there are reports of a shortening of the spine neck in potentiated spines during induction (Tonnesen et al., 2014) we explored both the effect of increasing and decreasing the length of the spine neck. Our results show that spine neck length plays a significant role in establishing the value of *L_crit_*. Figure 8C shows that as the length of the spine neck increases the value of Lcrit decreases slowly. Interestingly, a reduction of the spine neck length suggests that the inactive spine can remain isolated only at significantly larger distances.

Finally, the diameter of the spine neck seems to have a major effect in *L_crit_*. The diameter of the spine neck has been found to be regulated by neuronal and synaptic activity (Bloodgood and Sabatini, 2005; Tonnesen et al., 2014). Various experiments show a slowing of molecules passing through the spine neck as a result of high neuronal activity (Nishiyama and Yasuda, 2015), however there is evidence of the opposite effect as well (Tanaka et al., 2008; Tonnesen et al., 2014; Nishiyama and Yasuda, 2015). Our results are shown in Fig. 8D, and illustrate that a reduction of the spine neck diameter leads to a significant decrease in the critical distance between spines.

### Dependence of critical distance with activation characteristics of the molecular switch

The rate of protein synthesis in our model depends on the local protein concentration. This positive feedback mechanism is described through an activation function Θ(*c* − *c_θ_*). In all equations presented here, Θ is assumed to be a step-function. However, this approach is hardly realized in biological systems. Moreover, in our simulations we have replaced it by a steep Hill-function, with exponent *n* = 40 (see Table 1 and Eq. 9).

Here we explore the effect of relaxing the sharp steepness condition and determine the effect of decreasing the activation slope (i.e., decreasing *n*) in the critical distance *L_crit_*. We apply this to the case when polyribosomes are located in spine heads and assume that all active spines are still operating in the saturation regime. Consequently Eq. 4 is still valid except that the function Θ has been modified. The results are presented in Fig. 9.

**Figure 9:**
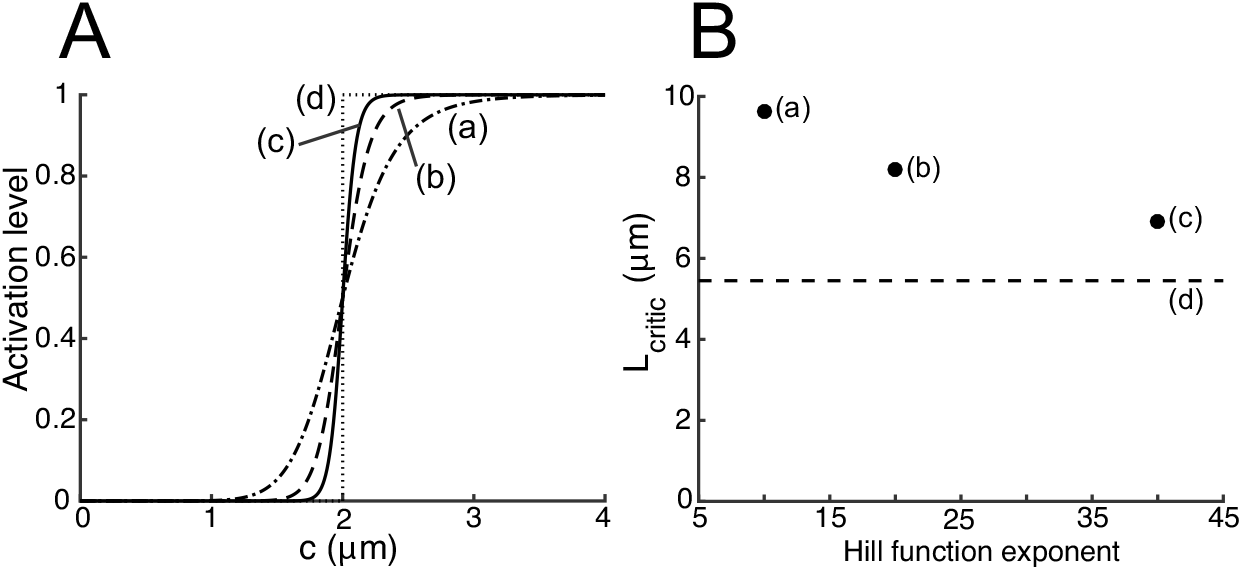
Dependency of critical distance with steepness of activation curve. A) Illustration of the different activation curves used in the simulations presented in panel B. Labels a, b and c correspond to those of panel B. B) Critical distance between spines increase as the steepness of the activation curve Θ decreases. Results were obtained for the standard parameters (see Table 1) and characteristic length constant λ= 120*μ*m. Points labeled a, b and c correspond to values of the exponent *n* in Eq. 9 equal to 10, 20 and 40, respectively. Point d, represents the result using a step-function as activation curve.

Panel B shows the calculated value of *L_crit_* as a function of the exponent *n* in Eq. 9. The circle marked as (c) corresponds to the standard parameters and the dashed line to the case of using a step-function as the activation curve. Panel A illustrates the changes in the shape of the function Θ corresponding to different values of *n* in B. The results show a dependence with the slope of the activation curve. A decrease in the slope by a factor of 4, increases the critical distance by ~ 2.5 *μ*m. These changes are relatively small, showing that our results do not depend strongly on the steepness of the activation function.

### Confirmation of methodology using 3D simulations

All results shown here, both analytical and computational have been based on a 1D approximation. This approximation is valid if there is no significant variation in concentrations across the different compartments. More specifically if concentration gradients along the length of spine-head, spine-neck and dendrite are much larger than the gradients at orthogonal directions. To verify that our results do not arise from this approximation, or are significantly altered by it we simulated 3D compartmental models using NEURON. Because 3D simulations are very computationally extensive, these simulations we carried out for the case of *N* = 1. Here the center spine was in the down state, and the flanking spines in the up state. This simulation case carried out for *L* = 17 *μ*m, a bit more than *L_crit_* for these parameters.

In figure xxA we show the 3D results at steady state for one parameter set. In figure xxB and xxC we compare results of 1D and 3Ds. The 3D simulation results are averaged over the volume of the structure in order to provide a comparison to the 1D results. We find that the concentrations are very similar in the 1D (black line) and 3D cases (red line). The 3D case seems like a somewhat smoothed version of the 1D case. However, due to the similarity between the results in 1D and 3D our results are robust to the precise method used.

Previous results are

## Discussion

The maintenance of long-term changes in synaptic efficacies of activated synapses most likely requires the ongoing synthesis of specific synaptic proteins. Such proteins remain active for long periods of time and might be degraded slowly. These proteins will diffuse and potentially reach the site of other synapses not previously activated, thus compromising the specificity of the initial activation pattern. Here we explored, using computational and analytical methods, the conditions under which synapses can remain isolated from each other during the maintenance of the protein synthesis-dependent phase of synaptic potentiation or L-LTP.

In order to provide a measure of the isolation of a synapse, we calculated the minimum distance (*L_crit_*) between synapses that would leave an inactivated synapse in its down-state while all other synapses around it are in the up-state. Here down- and up-state refers to the level of protein concentration at the location of the synapse; the up-state refers to stable, sustained high levels of protein being synthesized as the result of expressing L-LTP; the down-state refers to a low or basal level of protein concentration, typical of inactive synapses.

We used a simple bi-stable switch to model the protein synthesis process comprised of a feed-forward and degradation mechanism. In our setting the location of a switch corresponded with the location of a synapse, whether activated or not, and these switches were immobile. The main quantity representing the combined effect of diffusion and degradation of proteins is the characteristic length constant λ (see Eq. 6 in Materials & Methods), which can be thought of as the distance over which the concentration decays to ~ 36% (i.e., 1/*e*) of its magnitude at the (point)source (see Eq. 7 in Materials & Methods).

Our main result is presented in Fig. 7B. Here we show a comparison of *L_crit_* as a function of the characteristic length constant λ for two scenarios of the location of protein synthesis: in dendritic shafts and dendritic spines. Our calculations clearly show that the synthesis of proteins inside dendritic spines is necessary for synapse isolation. Here distances of less than 1*μ*m can be achieved already for values of λ < 60*μ*m, equivalently to degradation times of the order of hours. In contrast for these same values of λ, the minimum distance obtained when protein synthesis occurs in dendritic shafts is ~ 100 *μ*m. Note that the observed distance between spines is on the order of a few *μ*m, and synapse specificity during maintenance has been experimentally observed at the scale of a few *μ*m (Govindarajan et al., 2011).

We conclude from this that in order for synapses to remain isolated from each other during the maintenance phase of memory, it is necessary for polyribosomes to be inside dendritic spines (cf.(Bourne et al., 2007)). These results also imply that in synapses that lack spines such as inhibitory synapses, this method of long-term maintenance can not provide the same measure of synapse specificity. On the basis of our results we predict that if activated synapses in dendritic spines express L-LTP, for instance by showing an increase in the level of *PKMζ* (Palida et al., 2015) or other relevant proteins, then these spines should also contain polyribosomes.

The location of protein synthesis within synaptic spines is necessary, but is not sufficient for obtaining small enough values of *L_crit_*. We also find that smaller values of λ are necessary. Smaller values of λ correspond to faster degradation rates or slower diffusion constants.

Slowly degrading proteins like *PKM*ζ or PKCλ, which have degradation times of at least 4 hours (Palida et al., 2015) or longer have values of λ ~ 120 μm (assuming a diffusion constant as in Table 1). This presents a challenge for these proteins since the critical distances would be relatively large (see Fig. 7B) and thus would not offer good isolation. However, if the activated form of the protein changes faster than its degradation or diffusion rate (cf. (Harvey et al., 2008; Lee et al., 2009)) then this could lead to a shorter λ. Because the activity of *PKM*ζ depends on its phosphorylation state (Jalil et al., 2015) it could be possible that if *PKMζ* is dephosphorylated this might cause a reduction in its active lifetime (i.e., larger K) and thus reduce the value of λ. Thus our results would indicate that for slowly degrading proteins involved in L-LTP, like *PKMζ*, to also satisfy conditions for synapse isolation, their active form should be found primarily in activated dendritic spines and not in dendritic shafts. Our previous single compartment model of PKMζ-dependent maintenance was more complex than the very simple switch used here (Jalil et al., 2015). Our model, on the basis of experiemtnal observations, included two phosphorylation sites on the *PKM*ζ protein that control its activity. In such a model the relevant length scale is of the active-phosphorylated form of the protein, which is likely to have a shorter length-scale than total protein concentration.

Here we also explored how diffusional and morphological characteristics of the dendritic spine might improve conditions for synaptic isolation. Our main result is shown in Figures. 8C and especially 8D, which illustrate that there is a significant decrease in *L_crit_* as the diameter of the spine neck becomes smaller. This suggests that a further condition necessary for synaptic isolation would be that synapses at dendritic spines undergoing L-LTP would have narrower spine necks. However, this condition poses a problem as this might lead to making the synapse electrotonically isolated.

This condition is further supported by experimental evidence showing that the key regulatory component of the dendritic spine is the cross-section of the spine neck and examples of spines with high restrictive necks have been observed (Bloodgood and Sabatini, 2005; Tonnesen et al., 2014). Although changes in spine neck diameter have been reported to occur also in the opposite direction during induction (Tonnesen et al., 2014), it has not been verified experimentally if this trend remains or reverses during L-LTP.

Synaptic specificity—the property of synapses whereby only activated synapses undergo modifications in their synaptic efficacies while neighboring synapses remain unaltered—has been verified experimentally during LTP induction (Andersen et al., 1977; Lynch et al., 1977; Harvey et al., 2008) most results are confined to the early phases of LTP. Results presented in Govindarajan et al. (2011) indicate that synapse specificity in preserved for spines undergoing an L-LTP induction protocol and that such synapse specificity persists for many hours. In addition these results show that synapses in which L-LTP was induced, are capable of facilitating L-LTP in nearby spines stimulated 40 min later even when the second one is stimulated in the presence of protein synthesis inhibitors. This phenomenon called synaptic tagging and capture was previously demonstrated (Frey and Morris, 1997) using field potentials, these results use more precise techniques to delineate the limited spatial and temporal extent of this phenomenon. An interpretation of these results suggests that that proteins synthesized in one spine can diffuse into a second one up to distances of approximately 70 *μ*m; the kinetics and nature of these proteins are not yet identified. These results indicate that although long-term plasticity is localized to a synaptic spine, other aspects of protein synthesis depend plasticity are less local. However, given that we do not know what these processes are, it is difficult to model them in a mechanistic way.

Here is has been assumed that synapse specificity is necessary and desirable. It has been suggested that there are advantages to having less synapse specificity (Govindarajan et al., 2006),. Less specificity could generate clusters of strong synapses, which in turn could recruit dendritic spikes and can be used for local dendritic computations. However, here we find that if spine spacing is uniformly shorter than *L_crit_*, whole dendritic branches will potentiate. In order to generate clusters of limited extent there must a be highly non-uniform distributions of spines, with gaps larger than *L_crit_*, or there must be some diffusion barriers within the dendrites.

There is strong experimental evidence to support both the ideas of synaptic specificity and the memory-maintenance role of proteins. In this work we have addressed the question of how the diffusive nature of the latter can impose limits on the former. We propose here that for these two features of synaptic plasticity to co-exist, dendritic spines expressing L-LTP must contain polyribosomes, express only the active form of key proteins (e.g., de-phosporylated form of *PKMζ*) and have small spine neck diameters.

## Acknowledgements

This work is supported by NIH funding RO1 DA034979 (H.Z.S., T.C.S.), RO1 MH53576 (T.C.S.), 2R37 MH057068 (T.C.S.), and the Light-Fighter Trust (T.C.S.).

